# Substantial seagrass blue carbon pools in the southwestern Baltic Sea are spatially heterogeneous, mostly autochthonous, and include relics of terrestrial peatlands

**DOI:** 10.1101/2022.05.19.492657

**Authors:** Angela Stevenson, Tadhg C. Ó Corcora, Wolfgang Hukriede, Philipp R. Schubert, Thorsten B.H. Reusch

## Abstract

Seagrass meadows have a disproportionally high organic carbon (C_org_) storage potential (‘blue carbon’) within their sediments and thus can play an important role in climate change mitigation via their conservation and restoration. However, high spatial heterogeneity is observed in C_org_, with wide differences seen globally (i.e. tropical vs temperate), regionally, and even locally (within a seagrass meadow). Consequently, it is difficult to determine their contributions to the national remaining carbon dioxide (CO_2_) budget without introducing a large degree of uncertainty.

In order to address this spatial heterogeneity, we sampled 20 locations across the Baltic Sea coast of Germany to quantify carbon stocks and sources in *Zostera marina* seagrass-vegetated and adjacent unvegetated sediments. To predict and integrate the C_org_ inventory in space, we measured the physical (seawater depth, sediment grain size, current velocity at the seafloor, anthropogenic inputs) and biological (seagrass complexity) environment to determine regional (between sites) and local (within site) drivers of C_org_ variation.

Here we show that seagrass meadows in the German Baltic Sea constitute a significant C_org_ stock, storing on average 7,785 ± 679 g C/m^2^, 13 times greater than meadows from other parts of the Baltic Sea (outside of Germany), and four-fold richer than adjacent unvegetated sediments. Stocks were highly heterogenous; they differed widely between (by 10-fold) and even within (by 3 to 55-fold) sites. At a regional scale (350 km), C_org_ was controlled by seagrass complexity, fine sediment fraction, and seawater depth. Autochthonous material (seagrass-derived and large infauna) contributed to 78% of the total C_org_ in vegetated sediments and the remaining 22% originated from allochthonous sources (phytoplankton, drift algae *Pilayella littoralis*, and other macroalgae). However, relic terrestrial peatland material, deposited during the last deglaciation 5,806 and 5,095 years BP, was an unexpected and significant source of C_org_.

Collectively, German seagrass meadows in the Baltic Sea are preventing 8.14 Mt of future CO_2_ emissions. Because C_org_ is mostly produced on site, and not imported from outside the boundaries of the meadow, the richness of this pool may be contingent on seagrass habitat health. Disturbance of this C_org_ stock could act as a source of CO_2_ emissions. However, the high spatial heterogeneity seen across the region warrant site-specific investigations to obtain accurate estimates of blue carbon, and a need to consider millennial timescale deposits of C_org_ beneath seagrass meadows in Germany and potentially other parts of the southwestern Baltic Sea.

## Introduction

The oceans have absorbed approximately one-third of anthropogenic carbon dioxide (CO_2_) emissions to date (Mcleod et al. 2011), making the marine biome one of the largest carbon stores on Earth and thus an integral part of the climate change mitigation strategy. Despite having a relatively small global extent (0.5% of total ocean seafloor; Macreadie et al. 2021), coastal vegetated ecosystems, like seagrass meadows, mangrove forests, and tidal salt marshes, account for almost half of the total organic carbon (C_org_) buried in marine sediments (Duarte et al. 2005, Mcleod et al. 2011, Krause-Jensen and Duarte 2016), collectively estimated to mitigate approx. 3% of global CO_2_ emissions (Macreadie et al. 2021). Seagrass meadows alone have been estimated to contribute to 10% of the total buried C_org_ in ocean sediments (Duarte et al. 2005). Extraordinary rates of C_org_ accumulation and long-term storage in seagrass meadows have been attributed to high primary productivity (Hendricks et al. 2008, Duarte et al. 2013, Macreadie et al. 2014), efficient ability to capture particles from outside meadow boundaries (Kennedy et al. 2010, Miyajima et al. 2015, Oreska et al. 2018), a heavy network of roots and rhizomes that stabilize sediments and the carbon accumulated within them (Duarte et al. 2013), and formation of muddy anoxic sediments that prevent decomposition of the C_org_, which can thus be stored for centuries to millenia (Fourqurean et al. 2012, Duarte et al. 2013, Greiner et al. 2016).

Seagrass meadows are found in tropical and temperate bioregions, on the coasts of all continents (except Antarctica) and thus occur within the exclusive economic zones (EEZ) of many coastal nations, including Germany (Hemminga and Duarte. 2000, Short et al. 2007). Through careful management of seagrass habitats, such nations can use this natural carbon sink as a way to sequester part of their CO_2_ emissions. However, high spatial heterogeneity is observed in soil C_org_ storage potential, with wide differences seen globally (i.e. tropical vs. temperate environments), regionally, and even locally (within a seagrass meadow) (e.g. Kennedy et al. 2010, Röhr et al. 2018, Prentice et al. 2020, Mazarrasa et al. 2021). Consequently, it is difficult to apply ecological economic approaches to provide accurate economic valuations and determine seagrass habitat contributions to the national remaining CO_2_ budget without introducing a large degree of uncertainty.

Regional estimates are contingent on site-specific evaluations because the mechanism involved in C_org_ storage is largely based on local environmental factors, such as (1) local hydrodynamic regimes (e.g. seawater depth, Lavery et al 2013, Serrano et al. 2014, Mazarrasa et al. 2017a; decreased water motion, Prentice et al. 2019; lower wave height and exposure, Samper-Villarreal et al. 2016), (2) anthropogenic inputs (Macreadie et al. 2012, Mazarrasa et al. 2017a,b, Ricart et al. 2020), (3) seagrass properties (e.g. species composition, Serrano et al. 2014, 2019; increased seagrass complexity, Jankowska et al. 2016, Samper-Villarreal et al. 2016, Mazarrasa et al. 2018, 2021), and (4) sediment characteristics (e.g. Dahl et al. 2016, Röhr et al. 2016, Gullström et al. 2018, Miyajima et al. 2017).

The Baltic Sea coast of Germany is home to lush *Zostera marina* seagrass meadows, where seagrasses span a total area of approximately 285 km^2^ between 1-8 m seawater depth (Schubert and Steinhardt 2014, Schubert et al. 2015, Schubert and Schygulla 2016). Provided that the ambitious nutrient abatement targets of the Baltic Sea Action Plan are met, there is the potential that seagrass meadows could expand by at least 57 km^2^ by the year 2066 (Bobsien et al. 2021) and restoration activities could increase the existing area, leaving a large potential to gain negative emissions via restoration or improved growing conditions of these habitats

The objectives of the present study were to (i) provide a detailed assessment of the regional (between sites) heterogeneity of blue carbon stocks along 350 km of German Baltic Sea coastline; (ii) compare C_org_ content between seagrass-vegetated and adjacent unvegetated sediments to understand local (within site) C_org_ variation; (iii) determine the source of C_org_ contributing to these stocks (autochthonous vs allochthonous); (iv) combine biophysical parameters such as seawater depth, sediment grain size, current velocity at the seafloor, and seagrass complexity with C_org_ content into a predictive model to understand regional drivers of C_org_ variation; (v) scale up measurements and convert to CO_2_ equivalent units to determine the role of seagrass conservation in the total CO_2_ budget of Germany.

## Materials and methods

### Study area

Sampling took place in 20 seagrass meadows in the western part of the Baltic Sea, along the coasts of Schleswig-Holstein (n = 17) and Mecklenburg-Vorpommern (n = 3) in northern Germany (Fig. 1). The German Baltic Sea coast consists of shallow bays and fjords that experience weak water currents and low wave heights (Petterson et al. 2018). Like the whole Baltic Sea region, German coastal waters were geologically shaped by the last glacial periods (Schmölcke et al. 2006, Andrén 2012). As a consequence of the last deglaciation, a conglomerate of differently sized stones, sand, and clay settled in the southwestern part of the Baltic Sea basin. The seabed consists of shallow sandy and muddy layers with consolidated marl underneath. A maximum water depth of 40 m is reached in the western part of the Baltic Sea. However, seagrasses here are rarely observed deeper than 8 m seawater depth (Schubert et al. 2015). The Baltic Sea is the largest brackish water basin in the world and because of the narrow Danish Straits connecting the Baltic Sea to the North Sea, low rates of water exchange are observed (residence time of 35-40 years) resulting in high eutrophication from nutrient discharge by the nine Baltic Sea nations that enclose it (HELCOM 2018). However, eutrophication, which negatively impacts seagrass health and bathymetric range, is most pronounced in Germany, Russia and Poland (Thorsøe et al. 2022).

**Figure 1.**
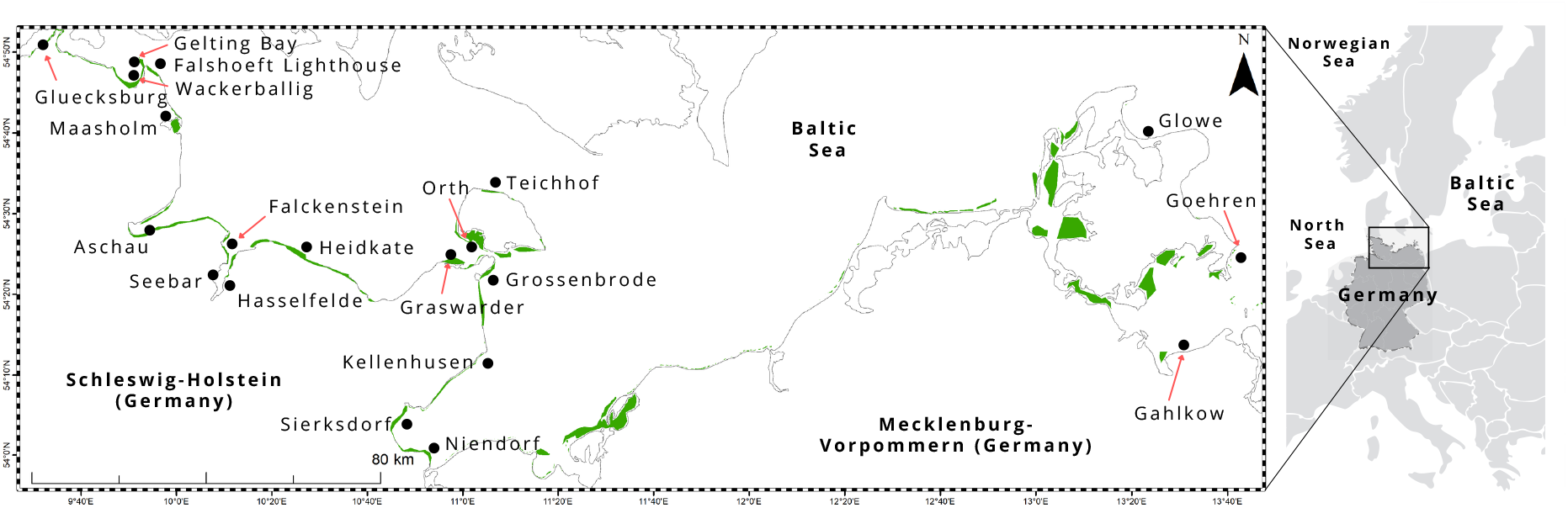
Study area, including 20 sampling locations, in the Baltic Sea coast of northern Germany. Seagrass area (green) was extracted from Schubert and Steinhardt 2014, Schubert et al. 2015, Schubert and Schygulla 2016.

Sites in the study area represent the greatest environmental gradient, ranging from wave exposed (e.g. Heidkate, Falshoeft lighthouse, Teichhof, Goehren) to sheltered (e.g. Orth, Maasholm), relatively pristine (nature reserve e.g. Gelting Bay and Graswarder; Aschau is protected from human impact due to military controlled access to the area) to varying degrees of anthropogenic inputs (adjacent to a marina e.g. Gluecksburg, Glowe; heavy ship traffic and near urban areas e.g. Kiel Fjord, including Falckenstein, Seebar, Hasselfelde; agricultural land e.g. Wackerballig; tourist areas e.g. Grossenbrode, Kellenhusen, Sierksdorf), and proximity to river influx (e.g. Gahlkow, Hasselfelde, Niendorf).

### Sample collection and core subsampling

A total of 169 sediment cores (30 cm length; 5.5 cm inner diameter) were sampled in seagrass vegetated meadows (n = 110 cores) and nearby unvegetated sediments (n = 59 cores). Nine cores (with some exceptions) were collected at each site, from three sublocations: (1) in the high-density part of the meadow (‘dense sublocation’ hereafter), (2) low density or fringe of the meadow (‘sparse sublocation’ hereafter), and (3) an adjacent unvegetated sediments at least 5 m from seagrass. Dense and sparse sublocations are collectively known as seagrass-vegetated sediments. Seagrass sediment cores were sampled at least 10 m apart from each other. Seawater depth was measured from a diver computer. Seagrass shoot density was counted using a 0.04 m^2^ frame, and the leaf lengths of five randomly selected plants were measured next to the location where the core was extracted. Seagrass density and leaf length vary between sites and at meadow stage of maturity. The measure of “seagrass complexity” was defined as the product of seagrass canopy height and shoot density, to obtain the sum of leaf heights within a unit of area (in m/m^2^) (Prentice et al. 2019).

Cores were collected between 1-5 m seawater depth manually via SCUBA divers (self-contained underwater breathing apparatus) pounding impact resistant PVC tubes into the sediment with a rubber mallet. Cores were capped at both ends by the divers and stored upright for transport to shore, and in a cooler thereafter for transport to the lab. Cores were stored at 0°C until further processing.

Sediments are known to shift during the coring process, so the degree of compression was measured by divers once the core was fully inserted. Compaction was calculated as the distance (in cm) from the top of the core to the sediment surface outside of the core, divided by the sample depth (cm of compression/cm core depth). A compression correction factor was calculated by dividing the length of the sample recovered by the length of core penetration (Howard et al, 2014). Cores were subsectioned into 5 cm depth intervals, and depth intervals were adjusted based on compaction.

Sediments from each depth interval were homogenized, measured for dry-bulk density and subsampled for chemical analyses (total C_org_, δ^13^C, δ^15^N, ^14^C, each described in detail in subsections below). A total of 394 sections underwent total C_org_ analyses: all top (0-5 cm) layers of all cores were processed for total C_org_, but the selection of subsequent core depth intervals depended on color changes throughout the core (color changes indicate potential changes in C_org_ content). If no color change was apparent, the mid (10-15 cm) and bottom sections of the core were processed (typically 20-25 cm, except where cores were shorter due to marl or difficult sediments). If a color change occurred, the respective section was instead examined. The 5-10 cm depth interval of all cores was used for grain size analysis (described below, under ‘grain size analysis’). Visible plant material (seagrass roots and rhizomes) and infauna (lug worm *Arenicola marina*, and, in some cases, soft tissue of clams and mussels) were removed from the sediment prior to chemical analyses (hereafter referred to as ‘visible organic’), but all other materials, such as wood and shells, were left in the sediment (referred to as ‘particulate organic carbon’ or ‘POC’).

### Maximum orbital velocity

These values were represented as maximum wave-generated orbital velocity (MOV) at the seafloor, modelled for the year 2021. Due to the limited availability of simulated wave data, MOVs were calculated only for cores taken at the outer coast sites of Schleswig-Holstein, not for the Schlei fjord site (Maasholm) and also not for any of the sites in Mecklenburg-Vorpommern (Gahlkow, Glowe, Goehren). MOV was calculated as a function of wave height, mean wave period, mean wave length (all simulated values), and seawater depth for each site according to linear wave theory as per the procedure described in Bobsien et al. (2021).

### Grain size distribution

The 5-10 cm depth interval of each core was processed for grain size distribution. Visible organics were first removed, then the sediments were oven dried at 60°C for a minimum of 48 hrs. Dry sieving was conducted with stainless steel Test Sieve ISO 3310-1 and sieve shaker (Fritsch Analysette 3 Spartan Pulverisette 0) set to amplitude 2 mm for 15 min. The fractions of sediment in 2 mm, 1 mm, 500 μm, 250 μm, 125 μm, 63 μm, and <63 μm (herein referred to as ‘fine sediment’ fraction) size classes were determined to the nearest 0.01 g, and the percent amount of each size class was calculated using the total sample weight obtained from the sum of each fraction.

### C_org_ stock quantification

Sediment total C_org_ was determined using an Elemental Analyzer (EURO EA Elemental Analyzer). Sediments were first dried at 60°C for 48 h and then ground to a homogeneous fine powder using a mechanical agate ball mill (Fritsch Pulveisette 5) set at rotational speed 240-300 rpm for 15-20 minutes. A subsample of this homogenized sediment was acidified to remove inorganic carbon by adding 1 M HCl drop-by-drop until gas evolution ceased. Samples were observed under a dissecting microscope to ensure CO_2_ had fully evolved. These acidified samples were re-dried at 60°C for 48 h, ground again (with mortar and pestle), and encapsulated into silver capsules. Total organic carbon concentrations were calculated based on a linear regression. Acetanilide and sediments were used as standards to measure data accuracy and analytical uncertainty.

### Stable isotope analyses

To decipher the source material of the remaining (non-visible) organic fraction in the sediments (referred to as POC), four sites (Falckenstein, Graswarder, Grossenbrode, Wackerballig) underwent further examination using biotracers of δ^13^C and δ^15^N. Samples were pre-treated as outlined above (see ‘C_org_ stock quantification’), and then combusted in an elemental analyzer system (NA 1110, Thermo) coupled to a temperature-controlled gas chromatography oven (SRI 9300, SRI Instruments), connected to an isotope ratio mass spectrometer (DeltaPlus Advantage, Thermo Fisher Scientific) as described by Hansen et al. (2009). δ^15^N, δ^13^C ratios were reported in delta notation relative to the international Air-N2 and VPDB scale following the equation:

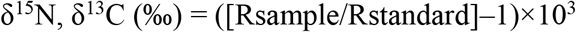

where R =^15^N/^14^N or ^13^C/^12^C. N_2_ and CO_2_ gases were used as reference gases and calibrated against International Atomic Energy Agency (IAEA) reference standards (N1-,N2-, NO3-) and National Institute of Standards and Technology (NBS-22 and NBS-600) compounds. To measure analytical uncertainty, acetanilide and caffeine were used as internal standards after every sixth sample to test if the analytical setup was working properly. Precision for Caffein was ± 0.13 for ‰ δ^15^N and ± 0.09 ‰ for δ^13^C; and ± 0.20 ‰ for δ^15^N and ± 0.27 ‰ for δ^13^C for Acetanilide.

A two biotracer (δ^13^C and δ^15^N), five-source Bayesian mixing model using R package (R Core Team 2019) MixSIAR (Stock and Semmens 2016) was used to determine the contribution of C_org_ to seagrass-vegetated and unvegetated sediments in Falckenstein, Großenbrode, Graswarder, and Wackerballig. Signatures of five potential sources were compiled from previous studies in Kiel Fjord, Germany and Gulf of Gdansk, Poland, and consisted of: (1) *Pilayella littoralis*, a filamentous brown algae known to form thick drifting mats accumulating in seagrass meadows in Germany (Kruk-Dowgiallo 1991, Kiirikki and Lehvo 1997), (2) other macroalgae (e.g. *Ulva intestinalis* and *Cladophora fracta*) extracted from Table 2 in Maksymowska et al. 2000; and (3) phytoplankton (collected by seston), (4) epiphytes attached to seagrass leaves, (5) seagrass leaves (there is little difference between above- and below-ground biomass signatures, see Röhr et al. 2016) extracted from Table 1 in Mittermayr et al. (2014). Sampling location (four locations) and vegetation state (two states: vegetated vs unvegetated) were included as random effects in the stable isotope mixing model. Vegetation state was nested within sampling location for the random structure. We specified an uninformative, generalist prior and discrimination factor of 0 as no further fractionation is expected in stored C_org_ (Greiner et al. 2016, Miyajima et al. 2017).

**Table 1.** Results of the Generalized Linear Mixed Model (Gamma, log link function), testing for the effect of sublocation on local (within site) sediment organic carbon (C_org_) content of seagrass meadows in the German Baltic Sea. ‘Dense’ sublocation refers to the high-density part of the meadow, ‘sparse’ sublocation is the low density or fringe of the meadow, and ‘unvegetated’ refers to adjacent unvegetated sediments at least 5 m from seagrass. Statistically significant effects in bold; α= 0.05.

| Source of error | Estimate | Std. Error | t-value | p-value |
| --- | --- | --- | --- | --- |
| Intercept | 8.9842 | 0.2465 | 36.499 | **<0.0001** |
| Sparse location | -0.7258 | 0.3492 | -2.078 | **0.04** |
| Unvegetated sediments | -2.5450 | 0.3470 | -7.335 | **<0.0001** |
| **Posthoc test** |  |  |  |  |
| Sparse vs Dense | -0.7258 | 0.3492 | -2.078 | 0.09 |
| Unvegetated vs Dense | -2.5450 | 0.3470 | -7.335 | **<0.001** |
| Unvegetated vs Sparse | -1.8192 | 0.3475 | -5.234 | **<0.001** |

**Table 2.** Radiocarbon ages (in years before present, BP) of wood material collected from *Zostera marina* seagrass-vegetated sediment cores in Sierksdorf, Luebeck Bay, in northern Germany. In some instances, two different wood pieces were dated from the same core section. Note: the material originated from two distinct time intervals (avg. 5,806 and 5,095 BP; younger time interval in bold), with a hiatus of approx. 700 years. BP is related to the hypothetical atmospheric value of 1950; SD - standard deviation.

| Core ID | Core depth interval<br>(cm) | $^{14}\text{C}$ age<br>(Year BP ± SD) |
| --- | --- | --- |
| C2 | 5-10 | 5,755 ± 30 |
| | 15-20 | 5,795 ± 30 |
|  | 15-20 | **5,131 ± 29** |
| C4 | 5-10 | 5,850 ± 30 |
| | 15-20 | 5,920 ± 30 |
| | 20-24 | 5,775 ± 29 |
| | 20-24 | 5,745 ± 35 |
| C7 | 5-10 | **5,116 ± 28** |
|  | 5-10 | **5,044 ± 28** |
|  | 10-15 | **5,090 ± 28** |
| C12 | 5-10 | 5,805 ± 35 |

### Radiocarbon dating

Large amounts of exceptionally well-preserved wood pieces were discovered in sediment cores extracted from the seagrass meadow in Sierksdorf, Luebeck Bay (Fig. 2). Of those, eleven pieces from a depth of 5 to 24 cm in four sediment cores were radiocarbon dated. The samples were visually inspected under a microscope and an appropriate amount of wood material was selected for dating. A standard decontamination procedure was subsequently applied to remove carbonates and soil humic contaminants, consisting of 1% HCl, 1% NaOH at 60 °C and again 1% HCl. All radiocarbon measurements were conducted at the Leibniz-Labor, using the type *HVE 3MV Tandetron 4130* accelerator mass spectrometer (AMS). Following standard procedures, the ^14^C/^12^C and ^13^C/^12^C isotope ratios were simultaneously measured by AMS, compared to the CO_2_ measurement standards (oxalic acid II), and corrected for effects of exposure to foreign carbon during the sample pretreatment. The resulting ^14^C-content was corrected for isotope fractionation, related to the hypothetical atmospheric value of 1950, and reported in pMC (percent Modern Carbon). This value was used to calculate the radiocarbon age according to Stuiver and Polach (2013). The reported uncertainty of the ^14^C result takes into account the uncertainty of the measured ^14^C/^12^C ratios of sample and measurement standard, as well as the uncertainty of the fractionation correction and the uncertainty of the applied blank correction. The radiocarbon ages were translated to calendar ages using the software package OxCal4 (Ramsey and Lee 2013) and the Intcal20 dataset (Reimer et al. 2020).

**Figure 2.**
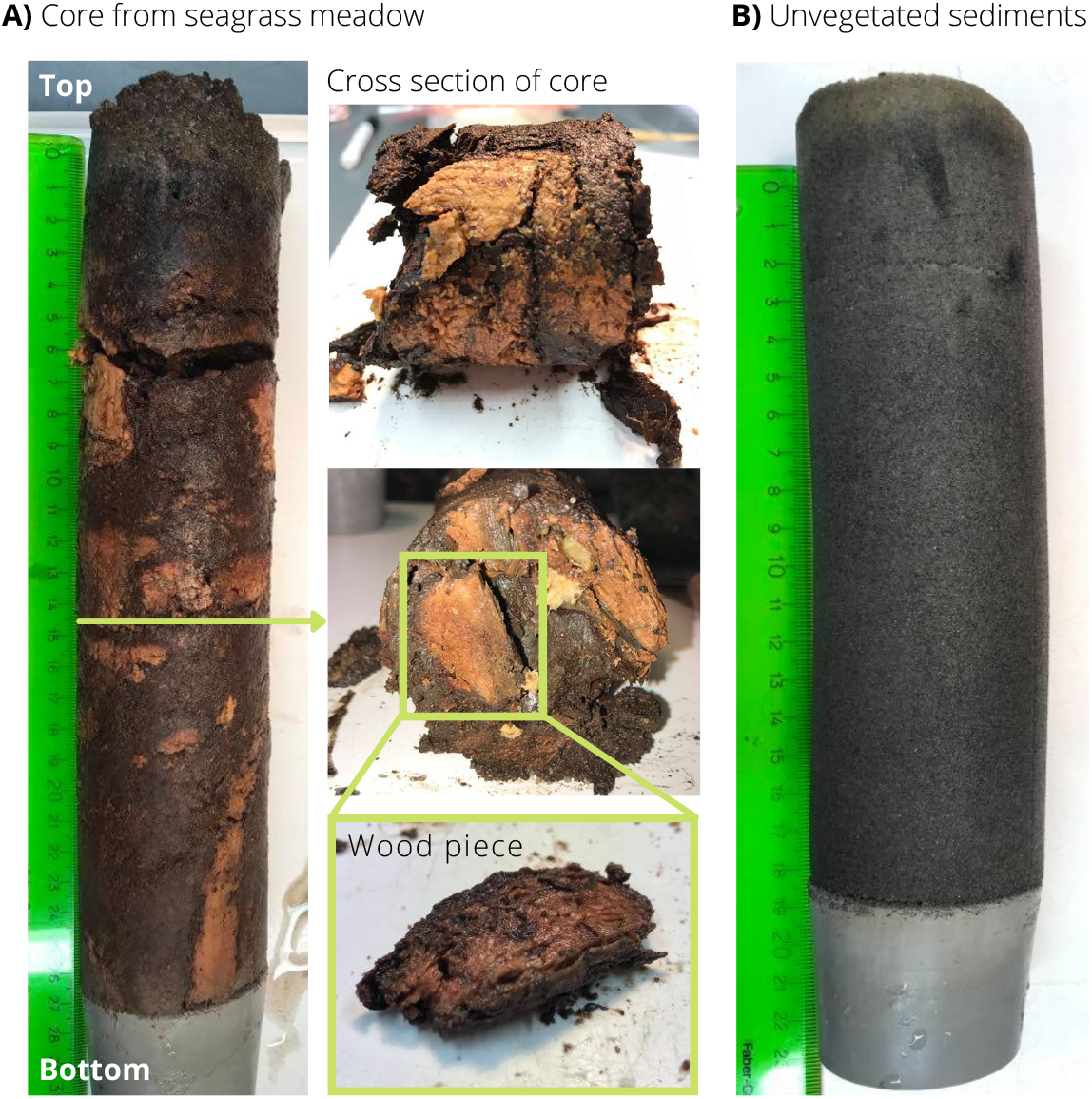
Core from *Zostera marina* seagrass-vegetated (A) and adjacent unvegetated (B) sediments in Sierksdorf, Luebeck Bay, where unexpected large amounts of exceptionally well-preserved wood pieces were discovered.

### Statistical analyses

Statistical analyses were performed in R version 4.1.3 (R Core Team 2022) for Mac OS X, and all mixed models were implemented with package “lme4” (Bates et al. 2015). For all tests, significance was determined at *p* < 0.05. Visual inspection of standard model validation graphs was used to verify model assumptions: residuals versus fitted values were used to verify homogeneity; a histogram or Quantile-Quantile (q-q) plot of the residuals for normality; and residuals versus each explanatory variable to check independence.

Generalized linear mixed models (GLMM) with a Gamma distribution and log link function were used to test: (1) core position effect on C_org_ between sublocations (dense, sparse, unvegetated sediments) within each site (locally), with a random intercept of sublocation nested in sampling location (site), and (2) biophysical factors influencing regional differences in stocks within seagrass-vegetated sediments, with sampling location as a random intercept. In the local model (1), a posthoc test using Tukey contrasts was performed with package “multcomp” (Torsten et al. 2008) to examine multiple comparisons between sublocations of each site. A multi-model inference approach based on AICc was used to determine the strongest predictors of regional C_org_ (2). Selection criteria AICc was used (instead of AIC) due to a small sample size (n/k < 40; n = total number of observations; k = total number of parameters in the most saturated model, including both fixed and random effects). The saturated model included four factors: seagrass complexity, average MOV, seawater depth, percent fine sediments, and their interactions. However, it is important to consider multicollinearity when interpreting model outputs as coefficient estimates (i.e. beta coefficients) and p-values become very sensitive to any small changes in the model. Multicollinearity of parameters in the dataset and among saturated model terms were examined using the Pearson correlation coefficient (*ρ*) and variance inflation factor (VIF). Correlation coefficients < 0.6 and VIFs approx. < 5 indicate an acceptable level of collinearity, VIFs > 10 warrant further investigation (Montgomery and Peck 1992). Low collinearity was observed in the data itself (*ρ* = −0.43 to 0. 35), but structural multicollinearity (between some model terms) was observed in the saturated model (VIF 1.876 to 14.051), so all terms with VIF >10 were removed from the global model. Hence, the final, global model included four factors: seagrass complexity, average MOV, seawater depth, percent fine sediments and two-way interactions between all variables except average MOV and fine sediment fraction, and all three- and four-way interactions were excluded (excluded VIFs: 10.055 to 14.051). After model selection, statistically indistinguishable (ΔAICc <2) candidate models were averaged using package ‘MuMIn’ (Bartoń 2022). To estimate the importance of each variable, the RVI (relative variable importance) value was computed by summing Akaike weights across all averaged models where a particular variable appeared. Model-averaged coefficients and p-values were obtained from the ‘full average’ (rather than the ‘conditional average’).

## Results

### C_org_ stocks

Averaged across all seagrass sediment cores, including dense and sparse sublocations (n = 110 cores total) and integrated to 25 cm core length, C_org_ stocks averaged at 7,785 + 679 g C/m^2^, and varied 10-fold between sites, ranging from 2,151 + 569 and 22,518 + 3,753 g C/m^2^ (Fig. 3). The highest average C_org_ stocks were found in Maasholm (Schlei Fjord) and Orth (Fehmarn Island), while the lowest values were observed in Falckenstein (Kiel Fjord) and Niendorf (Luebeck Bay).

**Fig. 3.**
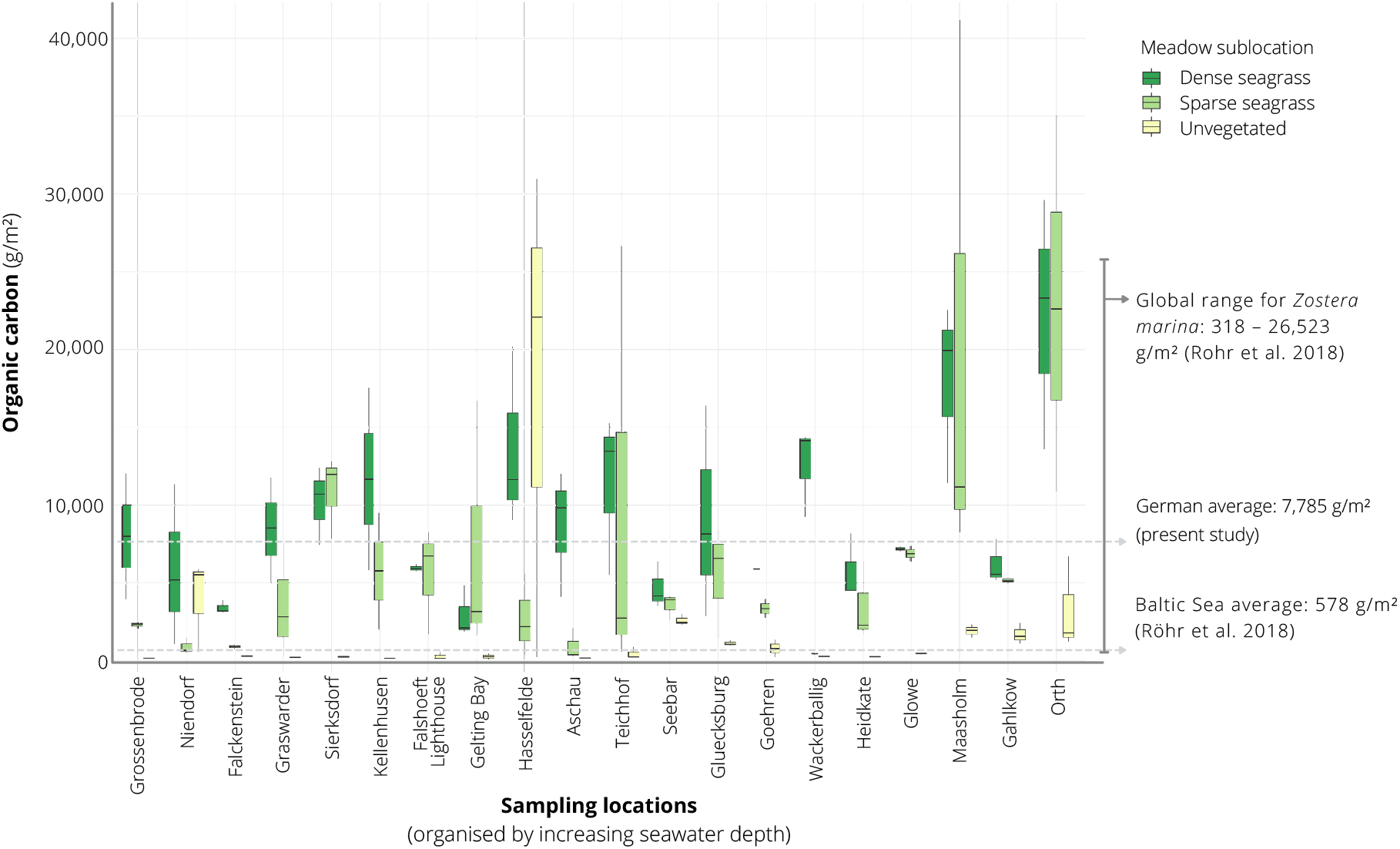
Overview of the organic carbon (C_org_) stocks (integrated to 25 cm sediment depth) across 20 sampling locations situated in the Baltic Sea coast of northern Germany. Sites are ordered by increasing seawater depth (left to right). Box and whiskers show the median as a line, first and third quartiles as hinges, and the highest and lowest values within 1.5 times the inter-quartile range as whiskers.

An examination of the local (within site) variation in C_org_ content across all sites showed significantly greater (by four-fold) stocks in seagrass-vegetated sediments (both dense and sparse sublocations) compared to nearby unvegetated sediments (dense: average 9,210 + 814 g C/m^2^; sparse: 6,359 + 1108 g C/m^2^; unvegetated: 1,840 + 666 g C/m^2^) (Table 1A, B; Fig. 3). However, considering sites individually, within site comparisons ranged widely: in all but three cases (Hasselfelde, Niendorf, Seebar), seagrass sediments had 3 to 55 times more C_org_ than nearby unvegetated sediments, with Kellenhusen, Sierksdorf, and Wackerballig showing the greatest discrepancies (>34 times that of unvegetated sediments). Gahlkow, Goehren, Gluecksburg, Maasholm, and Orth had the least difference (<10 times that of nearby unvegetated sediments). Niendorf and Seebar had the same, and Hasselfelde less C_org_ than nearby unvegetated sediments. C_org_ content did not differ significantly between sparse and dense sublocations.

### Sources of sediment C_org_ (δ^13^C, δ^15^N, ^14^C)

Overall, the stable isotope signatures of the POC fraction of seagrass-vegetated sediments (including dense and sparse sublocations along the entire core depth) were near identical to those of unvegetated sediments (δ^15^N 5.2 ± 0.3, ‰ δ^13^C-21.9 ± 0.2‰ vs δ^15^N 5.3 ± 0.2, δ^13^C-21.6 ± 0.2‰). In seagrass sediments, signatures ranged between δ^15^N 4.1 + 0.2 (Großenbrode) to 7.3 + 0.3 (Falckenstein), and δ^13^C-23.1 + 0.4 (Falckenstein) to −19.4 + 0.5 (Wackerballig) (Fig. 4B). δ^13^C and δ^15^N signatures were also homogenous across core depth; top sediments (0 to 5 cm) averaged across all seagrass cores were ^15^N depleted by 0.48‰ and ^13^C enriched by 0.68‰ compared to the 10 to 15 cm and 15 to 20 cm sections.

**Fig. 4.**
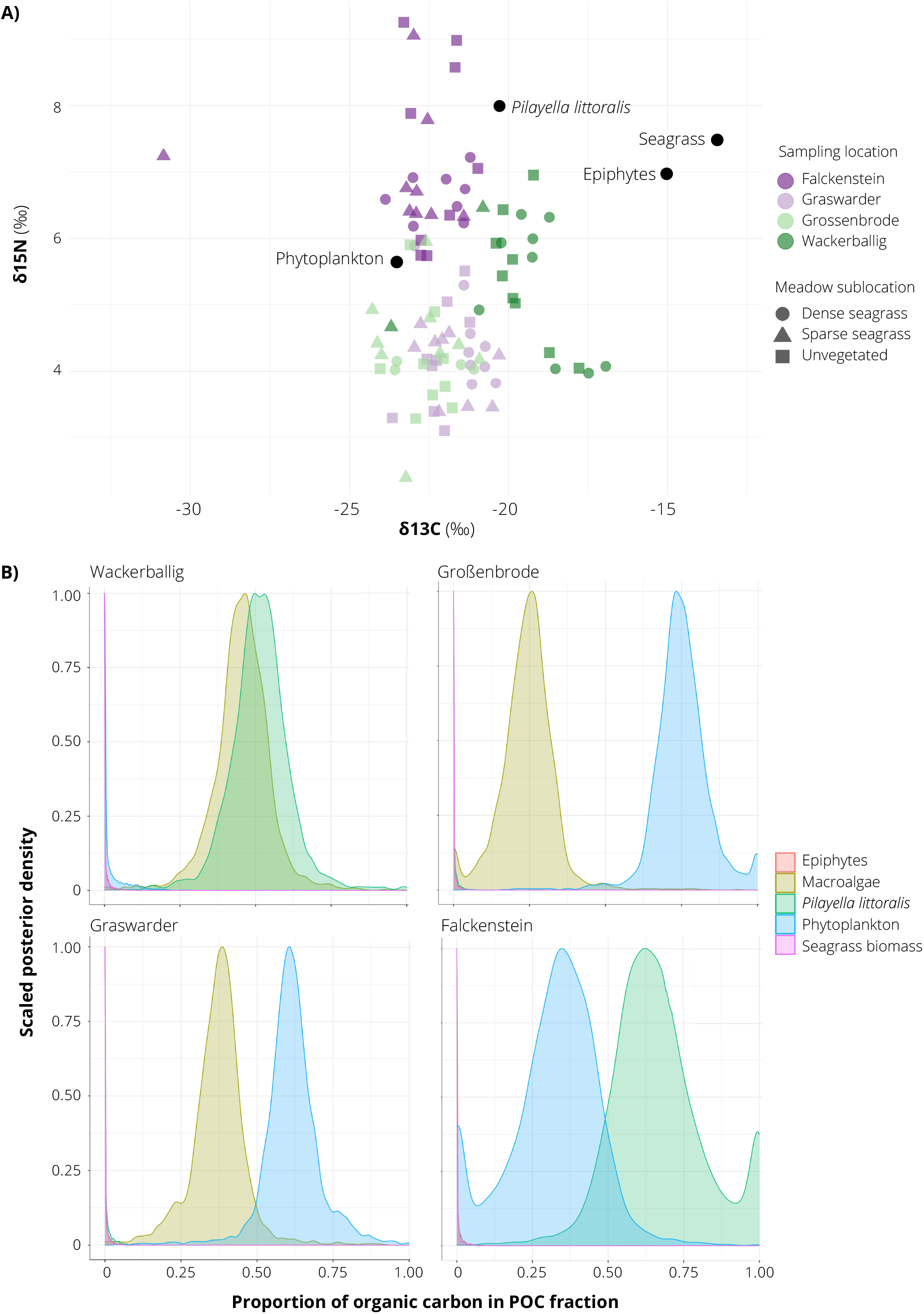
δ^13^ C, δ^15^N stable isotope signatures (A) and posterior estimates of the proportion of organic carbon (C_org_) sources (B) of sediments sampled in seagrass-vegetated and unvegetated sediments off the coasts of Falckenstein (FS), Wackerballig (WB), Großenbrode (GB), Graswarder (GW), in northern Germany. Sources (black circles) were obtained from Table 1 in Mittermayr et al. (2014) and Table 2 in Maksymowska et al. (2000).

Averaged across all sites, visible organic material such as invertebrates and seagrass root, shoot, rhizome material contributed 78% of the total C_org_ in seagrass-vegetated sediments and 29% in unvegetated sediments (excluding Hasselfelde, Niendorf, Seebar). Visible organics were responsible for >94% of the C_org_ in unvegetated sediments in Hasselfelde and Niendorf, but not in Seebar. Here, there were no visible organics, instead this material was exclusively of particulate origin (measured in the POC fraction). Results from the five-source two-biotracer (δ^13^C, δ^15^N) mixed model of the remaining (22%) organic fraction (the non-visible fraction, called POC) in the sediments of Falckenstein, Wackerballig, Großenbrode, Graswarder, showed that phytoplankton (24%), *P. littoralis* (18%) and other macroalgae (22%), made the largest contribution to C_org_ overall, not seagrass biomass or its epiphytes, and a combination of two of these three dominant carbon sources could be seen at each site (Fig. 4A). Sampling location accounted for most of the variation in the mixing model (50^th^ percentile *σ*= 6.311), while vegetation coverage (vegetated vs unvegetated) had a negligible effect (50^th^ percentile *σ*= 0.105).

Radiocarbon dating of wood pieces found in Sierksdorf cores revealed no age-depth correlation within a single core, but all dates fell in two well separated time intervals, averaging at 5,806 years BP and 5,095 years BP (before present), i.e. a hiatus of approx. 700 years between these time intervals (Table 2).

### Biophysical predictors of C_org_

The percent fine sediment fraction of seagrass-vegetated sediments varied greatly between sites, from 0.09 + 0.04% (Gahlkow) to 7 + 3% (Goehren). All but five sites had <2% fine sediments (Glowe, Goehren, Niendorf, Orth, Sierksdorf) (Table 3). Unvegetated sediments had a low fine sediment fraction (<3%), with Orth and Seebar ranking among the highest. Seagrass-vegetated sediments contained similar or more (by 1-31 times) fine-grained sediments than adjacent unvegetated sublocations, except in Graswarder, Gluecksburg, Seebar, where unvegetated sediments contained 2-10 times more fine-grained sediments than their vegetated counterparts. For seagrass complexity, Maasholm exhibited the lowest (92 + 21 m/m^2^) and Wackerballig the highest (539 + 132 m/m^2^) average seagrass complexity of all sites (including sparse and dense sublocations) (Table 3). Dense sublocations were one to five times more complex than sparse sublocations, with the greatest difference observed in Teichhof and Aschau, and smallest difference seen in Sierksdorf and Großenbrode. Average MOV varied six-fold, ranging from 0.155 + 0.007 m/s (Niendorf) and 0.91 + 0.03 m/s (Teichhof) between sties, but some (Falshoeft lighthouse, Heidkate, Kellenhusen, Teichhof, Wackerballig) experienced strong peak MOVs (> 3 m/s) (Table 3).

**Table 3.** Summary of average (± SE) of each biophysical variable calculated (seagrass complexity), modelled (maximum Maximum Orbital Velocity; MOV), or measured (seawater depth, % fine sediments) in 20 seagrass meadows along the Baltic Sea coast of Germany. Seagrass complexity was calculated by multiplying seagrass canopy height by shoot density measured next to each core, at each site. MOV was modelled only for outer coast sites in Schleswig-Holstein, thus excluding the inner waters of the Schlei fjord (Maasholm) and Rugen island (Gahlkow, Glowe, Goehren). ‘Average max. MOV’ - average maximum MOV value observed for each core location. ‘Fine sediments’ - percent fraction of the sediment with grain size <63 μm.

| Sampling location | Average seagrass complexity (m/m<sup>2</sup>) | Average MOV (m/s) | Average max. MOV (m/s) | Average seawater depth (m) | Fine sediments (<63 μm; %) |
| --- | --- | --- | --- | --- | --- |
| Aschau | $186 \pm 57$ | $0.5 \pm 0.2$ | $1.7 \pm 0.1$ | $2.7 \pm 0.3$ | $1.9 \pm 0.6$ |
| Falckenstein | $154 \pm 29$ | $0.184 \pm 0.001$ | $0.989 \pm 0.005$ | $3.7 \pm 0.0$ | $1.9 \pm 0.6$ |
| Gahlkow | $188 \pm 45$ | na | na | $1.12 \pm 0.05$ | $0.09 \pm 0.04$ |
| Gelting bay | $139 \pm 60$ | $0.360 \pm 0.005$ | $2.05 \pm 0.02$ | $3.0 \pm 0.0$ | $1.5 \pm 0.5$ |
| Falshoeft lighthouse | $224 \pm 39$ | $0.49 \pm 0.06$ | $3.5 \pm 0.3$ | $3.1 \pm 0.4$ | $0.4 \pm 0.1$ |
| Glowe | $293 \pm 64$ | na | na | $1.35 \pm 0.09$ | $4 \pm 1$ |
| Gluecksburg | $108 \pm 17$ | $0.32 \pm 0.02$ | $1.67 \pm 0.08$ | $2.1 \pm 0.2$ | $0.6 \pm 0.1$ |
| Goehren | $163 \pm 69$ | na | na | $2.03 \pm 0.03$ | $7 \pm 3$ |
| Graswarder | $241 \pm 47$ | $0.446 \pm 0.003$ | $2.071 \pm 0.008$ | $3.4 \pm 0.0$ | $0.16 \pm 0.06$ |
| Großenbrode | $210 \pm 30$ | $0.331 \pm 0.002$ | $1.959 \pm 0.008$ | $4.3 \pm 0.0$ | $0.4 \pm 0.1$ |
| Hasselfelde | $155 \pm 34$ | $0.3 \pm 0.1$ | $0.9 \pm 0.1$ | $2.7 \pm 0.5$ | $0.9 \pm 0.2$ |
| Heidkate | $268 \pm 60$ | $0.66 \pm 0.02$ | $3.32 \pm 0.07$ | $1.4 \pm 0.0$ | $0.24 \pm 0.07$ |
| Kellenhusen | $211 \pm 29$ | $0.45 \pm 0.04$ | $3.0 \pm 0.2$ | $3.1 \pm 0.3$ | $1.06 \pm 0.06$ |
| Maasholm | $92 \pm 21$ | na | na | $1.23 \pm 0.02$ | $1.2 \pm 0.2$ |
| Niendorf | $175 \pm 31$ | $0.155 \pm 0.007$ | $1.34 \pm 0.04$ | $4.2 \pm 0.2$ | $3.2 \pm 0.2$ |
| Orth | $239 \pm 66$ | $0.58 \pm 0.01$ | $1.36 \pm 0.03$ | $1.0 \pm 0.0$ | $2.7 \pm 0.7$ |
| Seebar | $100 \pm 22$ | $0.244 \pm 0.007$ | $1.00 \pm 0.03$ | $2.2 \pm 0.1$ | $0.8 \pm 0.2$ |
| Sierksdorf | $194 \pm 47$ | $0.265 \pm 0.003$ | $1.89 \pm 0.02$ | $3.3 \pm 0.0$ | $2.8 \pm 0.9$ |
| Teichhof | $246 \pm 73$ | $0.91 \pm 0.03$ | $4.3 \pm 0.1$ | $2.2 \pm 0.1$ | $1.3 \pm 0.5$ |
| Wackerballig | $539 \pm 132$ | $0.6 \pm 0.1$ | $3.3 \pm 0.5$ | $1.5 \pm 0.6$ | $2 \pm 1$ |

Six alternative candidate models were deemed statistically indistinguishable (ΔAICc < 2) from each other (summarized in Table 4A). Meaning, there was no strong support for one particular model. In the averaged model, seawater depth, seagrass complexity, and fine sediment fraction were similarly important in predicting C_org_ within seagrass-vegetated sediments (RVI 1.00 for each, Table 4B), and they had a significant negative (seawater depth) or positive (fine sediment fraction, seagrass complexity) effect on C_org_ stocks (see ‘Estimate’ in Table 4 B). Avg. MOV had a weaker (by approx. 2.5 times) and insignificant effect in predicting C_org_. There were no significant interactive effects between variables.

**Table 4.** Summary of alternative candidate Generalized Linear Mixed Models (Gamma, log link function) with ΔAICc < 2 (A) and their model-averaged coefficients (B) for biophysical predictors of regional sediment organic carbon (C_org_) content of seagrass-vegetated sediments along the Baltic Sea coast of Germany. In all models, location (site) was included as a random effect. Avg. MOV - average Maximum Orbital Velocity; SG - seagrass. Statistically significant effects of model-averaged coefficients (B) are in bold; α= 0.05. ‘+’ designates a main effect and ‘x’ an interaction.

| A. Summary of statistically indistinguishable candidate models |  |  |  |  |  |  |  |
| --- | --- | --- | --- | --- | --- | --- | --- |
| Candidate model | df | Log L |  | AICc | ΔAICc | Weight |  |
| Avg. MOV + fine sed. + SG complexity + depth + depth x SG complexity | 8 | -754.85 |  | 1527.78 | 0.00 | 0.24 |  |
| Avg. MOV + fine sed. + SG complexity + depth + fine sed. x depth | 8 | -754.94 |  | 1527.98 | 0.19 | 0.22 |  |
| Avg. MOV + fine sed. + SG complexity + depth + fine sed. x depth + avg. MOV x SG complexity | 8 | -753.91 |  | 1528.47 | 0.69 | 0.17 |  |
| fine sed. + SG complexity + depth | 6 | -757.70 |  | 1528.59 | 0.81 | 0.16 |  |
| Avg. MOV + fine sed. + SG complexity + depth + avg. MOV x SG complexity | 8 | -755.52 |  | 1529.12 | 1.34 | 0.12 |  |
| Avg. MOV + fine sed. + SG complexity + depth + avg. MOV x depth | 8 | -755.81 |  | 1529.70 | 1.92 | 0.09 |  |
| B. Model-averaged coefficients (full average) |  |  |  |  |  |  |  |
| Source of error | RVI | VIF | Estimate | Std. Error | Adj. SE | z value | p-value |
| Intercept | na | na | 8.78745 | 0.13957 | 0.14190 | 61.927 | <0.0001 |
| Avg. MOV | 0.84 | 1.623 | 0.11958 | 0.12247 | 0.12397 | 0.965 | 0.3 |
| Seawater depth | 1.00 | 1.386 | -0.29599 | 0.13372 | 0.13602 | 2.176 | 0.03 |
| Seagrass complexity | 1.00 | 2.008 | 0.27162 | 0.12118 | 0.12277 | 2.212 | 0.03 |
| Fine sediments | 1.00 | 1.138 | 0.27036 | 0.11570 | 0.11768 | 2.297 | 0.02 |
| Depth x SG complexity | 0.24 | 3.688 | 0.04696 | 0.09490 | 0.09527 | 0.493 | 0.6 |
| Depth x fine sed. | 0.39 | 1.423 | -0.08820 | 0.13271 | 0.13341 | 0.661 | 0.5 |
| Avg. MOV x SG complexity | 0.29 | 3.220 | -0.05116 | 0.10025 | 0.10089 | 0.507 | 0.6 |
| Avg. MOV x depth | 0.27 | 1.372 | 0.01990 | 0.07521 | 0.07562 | 0.263 | 0.8 |

## Discussion

### Spatial heterogeneity of blue carbon stocks in Germany (objectives i, ii)

C_org_ stocks in the top 25 cm of seagrass-vegetated sediments in the German Baltic Sea were high (average 7,785 + 679 g C/m^2^), and richer than adjacent unvegetated sediments, but differed widely between and even within sites. Regional heterogeneity observed here was comparable to previous regional evaluations of blue carbon in *Z. marina* meadows (e.g. Röhr et al. 2016; Prentice et al. 2019), but local heterogeneity was much greater, with previous measurements reporting <10 times more C_org_ content in vegetated vs unvegetated sediments (Table 5), or similar carbon content between these sublocations (Prentice et al. 2019, 2020; Mazarrasa et al. 2021; Krause et al. 2022). Overall, average C_org_ stocks in the German Baltic Sea fit within the global range reported for those of *Z. marina* (318 + 10 to 26,523 + 667 g C/m^2^), but more closely resembled *Z. marina* C_org_ stocks in the Mediterranean Sea (8,793 + 2,248 g C/m^2^), Funen area of Denmark, and Skagerrak coast of Sweden, than C_org_ stocks in other parts of the Baltic Sea (Table 5). In fact, German blue carbon stocks were on average much richer (by 13 times) than the Baltic Sea average (578 + 43 g C/m^2^, excluding Germany, Röhr et al. 2018). A similar geographical trend was also reported for sites along the coast of Sweden, where percent C_org_ in top sediments were 10 to 25 times higher along the west (in Skagerrak) vs south and east coasts (in the Baltic Sea) of Sweden (Jephson et al. 2008). Consistent with the present study, wide regional variation in C_org_ stocks were observed in eastern Jutland of Denmark, which included “carbon hot spots” (stocks as high as 26,523 + 667 g C/m^2^ in Thurøbund) that were comparable to those observed in Germany (22,518 + 3,753 g C/m^2^). Two biophysical parameters were thought to be responsible for the C_org_ hotspot found in Thurøbund: low wave exposure and high seagrass productivity (420 + 98 shoots/m^2^) - seagrass densities and exposures that were similar in magnitude to those measured in the sheltered bay of Orth (467 + 98 shoots/m^2^) where the highest C_org_ was found in Germany. It is worthwhile contemplating explanations for the higher C_org_ stocks reported in Skagerrak-Kattegat and southwestern Baltic Sea relative to other parts of this basin (Table 5). The rich geological history that shaped the Baltic Sea, including northward retreat of the Scandinavian ice sheet that caused land uplift in the south and changed coastal landscapes as a consequence of sea-level rise during the Littorina Transgression, but also shaped the geology that enhanced its ability to sequester C_org_. For instance, hard-bottom and rocky shores dominate in the northern coasts, whereas till material and sandy/muddy beaches are common in the south (Schiewer 2008).

**Table 5.** Spatial heterogeneity (average, min, max) of C_org_ (g C_org_/m^2^) in the upper 25 cm of *Zostera marina*-vegetated and unvegetated sediments of the Baltic Sea. Geographical positioning within Baltic Sea based on coastal morphology, sills, other topographical formations (Leppäranta and Myrberg 2009).

| Country<br>(number<br>of sites) | Geographical<br>division within<br>Baltic Sea | C<sub>org</sub> in *Z. marina*-vegetated<br>sediments (g C<sub>org</sub>/m<sup>2</sup>) |  | C<sub>org</sub> in<br>unvegetated<br>sediments (g<br>C<sub>org</sub>/m<sup>2</sup>) | Source |
| --- | --- | --- | --- | --- | --- |
|  |  | Mean | Range |  |  |
| Germany<br>(20) | Danish Straits<br>(SW Baltic Sea) | $7,785 \pm 679$ | $2,151 \pm 569$ to<br>$22,518 \pm 3,753$ | $1,840 \pm 666$ | Present study |
| Denmark<br>(Funen<br>region, 5) | Danish Straits<br>(SW Baltic Sea) | $6,005 \pm 1,127$ | 400 (estimate)<br>to $22,518 \pm$<br>$3,753$ | NA | Röhr et al. 2016 |
| Gullmar<br>Fjord,<br>Sweden | Skagerrak Strait | $3,500 \pm 410$ | NA | 500 (estimated<br>from Fig. 2) | Dahl et al. 2016 |
| Finland<br>(10) | Gulf of Bothnia<br>(N Baltic Sea) | $627 \pm 25$ | 400 to 1,300<br>(estimated from<br>Fig. 4) | NA | Röhr et al. 2016 |
| Askö,<br>Sweden | Gotland Sea (N<br>Baltic Sea) | $500 \pm 50$ | NA | 200 to 600<br>(estimated from<br>Fig. 2) | Dahl et al. 2016 |
| Poland (3) | Gotland Sea, (S<br>Baltic Sea) | $370 \pm 15$ | $125 \pm 6$ to $570 \pm$<br>$29$ | $134 \pm 4$ | Averaged from<br>Jankowska et al.<br>2016 |

### Sources of C_org_ (objective iii)

Our results suggest that the C_org_ accumulating in seagrass meadows in the German Baltic Sea is primarily (78%) originating from autochthonous sources, material already in the meadow itself, like seagrass plants and large infauna. Thus, most of the C_org_ making up stocks here are produced on site, and not imported from outside the boundaries of the meadow. Only a small fraction (22%) originated from allochthonous sources, and of the five C_org_ sources tested, this fraction was predominantly derived from a combination of phytoplankton, drift algae *P. littoralis*, and other macroalgae. These findings are consistent with those of neighboring Baltic Sea nations, like Denmark and Poland, where seagrass biomass was found to be the biggest contributor to sediment C_org_ pools (13 - 81%, Röhr et al. 2016; 40 - 45%, Jankowska et al. 2016). However, phytoplankton material was the primary source of C_org_ in Finland (43 - 86%), while seagrass made a relatively small contribution to the overall sediment C_org_ pool (1.5 - 32%) (Röhr et al. 2016), and along the northwest Pacific coast, *Z. marina* meadows primarily sequestered allochthonous C_org_, originating from plankton, terrestrial, and kelp sources rather than seagrass material (Prentice et al. 2019, Krause et al. 2022). The latter contributed less than 25.3% to the C_org_ pool. A country-wide analysis of blue carbon stocks in Australia revealed that coastal meadows in temperate regions were dominated by autochthonous C_org_ sources (72% was derived from seagrass), whereas allochthonous material prevailed in tropical meadows (64%) (Mazarrasa et al. 2021). In a global analysis (across 88 locations and multiple seagrass species) of seagrass blue carbon stocks, seagrass biomass accounted for approximately 50% of the C_org_ stocks, while the remainder originated from allochthonous sources (Kennedy et al. 2010). The primarily autochthonous source of C_org_ in Germany, and temperate bioregions in general, is interesting and might suggest that richness of the blue carbon pool could be contingent on the health of the seagrass habitat itself.

In the mixing model, sampling location (not vegetation coverage) was the main driver of the variation in C_org_ sources (for the non-visible fraction of C_org_; POC). Coastal landscape and differences in inputs to marine foodwebs, including currents, upwelling, as well as point (e.g. sewage outfall, rivers) and diffuse (e.g. atmospheric, rainwater runoff) sources of nutrients may help explain some of the dissimilarity observed between locations. For example, phytoplankton contribution was abundant in the seagrass meadow in Graswarder, but absent in Wackerballig. The former is situated near a point source contamination of sewage outflow (Schubert et al. 2013), and the latter is adjacent to a large marine national park. Phytoplankton composition and abundance are often used as bioindicators of enhanced nutrient concentrations (2000/60/EC, EU, 2000). For the drift algae *P. littoralis*, large mats can become entrapped in narrow fjords (Falckenstein) and bays (Wackerballig), whereas they may be carried away from sites along open coasts (Grasswarder, Großenbrode), which fits with the pattern seen in the sediment sources of the present study.

While C_org_ content was consistently higher in seagrass-vegetated vs unvegetated sublocations, two cases had similar (Niendorf and Seebar) or more (Hasselfelde) C_org_ in their unvegetated sediments. A similar phenomenon was previously reported for *Z. marina* meadows and other temperate seagrass species (see Prentice et al 2019, 2020; Mazarrasa et al. 2017a,b, 2021; Krause et al. 2022). Some of this discrepancy was attributed to the export of organic material originating from seagrass habitats to adjacent unvegetated sediments (Duarte and Krause-Jensen 2017). A similar spillover effect of C_org_ from adjacent seagrass meadows may be contributing to the high C_org_ accumulating in nearby unvegetated sediments at both sites since visible C_org_ (not POC, but also not seagrass biomass) was the main source of C_org_ in both vegetated and unvegetated sediments of Hasselfelde and Niendorf. For Niendorf, this could partially help explain low C_org_ stocks in vegetated sediments (second lowest in the study). In Hasselfelde, a dense community (several cm thick) of bivalve *Cerastoderma edule* populated bare sediments at this site (but not seagrass-vegetated sediments) and their soft tissue would have contributed to C_org_ sinks (or CO_2_ sinks) measured in these sediments. Autochthonous export is not a valid justification for site Seebar as bare sediments were dominated by non-visible C_org_ (100% POC) and these had a sludge-like consistency with the highest fine (muddy) particle fraction of all sites and sublocations in the study. Heavy anthropogenic pressure at this site may be contributing to the excess nutrient inputs that would elevate baseline C_org_ in the sediments (Nixon 1995, Short and Burdick 1996, Bowen and Valliela 2001, Nedwell et al. 2002). Also, this site was not populated by seagrass only 8 years prior to the study, which could have contributed to these findings as well (pers. obs. P. Schubert)

Unexpected large amounts of well-preserved wood pieces were found in one location: Sierksdorf, Luebeck Bay. Radiocarbon dating of this wood suggests that it was deposited here during two distinct time intervals, averaging at 5,806 BP and 5,095 BP, that coincide well with the second phase of the Littorina Transgression (dated approx. 6,000 to 3,800 BP) following the last deglaciation (Schmölcke et al. 2006, Kostecki et al. 2021). In this period, rapid changes occurred especially in the southwestern part of the Baltic Sea, where fast sea level rise combined with a flat landscape led to the widespread die off of forests situated along the coast. These events caused the demise of coastal alder woodlands that subsequently produced peat containing abundant alder-wood remains (Schmölcke et al. 2006). Indeed, the lack of age (^14^C) correlation across core depth is typical of natural wood pieces (rather than man made artefacts) because younger wood can be found in deeper parts of the core due to the roots of the living tree. The two distinct time intervals occurring 700 years apart may suggest a recolonization event took place at this site. Interestingly, the two time intervals are situated on either side of a brief cooling period (the Subboreal period), which started 5,650 BP and caused widespread natural environmental changes in the southwestern part of the Baltic Sea (Schmölcke et al. 2006).

The oldest known submarine peatlands globally are dated 5,616 + 46 years BP and correspond to the thick *matte* formed by *Posidonia oceanica*, a long-lived Mediterranean seagrass, which constitute deep and significant C_org_ stocks (Lo Iocano et al. 2008). *Z. marina* does not form similarly thick and old sedimentary deposits, but our investigation confirms that it too can store millennia of carbon by protecting former (and now submerged) forested peatlands, acting as a protective covering for these rich carbon deposits (Krause-Jensen et al. 2019). Even well-preserved prehistoric settlements have been previously discovered beneath *Z. marina* meadows in Denmark (Fischer 2011, Andersen 2013, Pedersen et al. 2017) and Germany (Goldhammer & Hartz 2017). Submerged wood artefacts are also present in the Baltic Sea coast of Germany, including a site in Neustadt near Sierksdorf (Klooß 2014), and they too overlap with seagrass habitats. Recognition that some seagrass meadows in Germany are sitting atop ancient terrestrial carbon deposits that are potentially several meters thick has important consequences for avoiding the re-emission of C_org_ stored on a millennial timescale. However, our understanding of submerged coastlines is incomplete, so it is likely that many more submarine peatlands, like that found in Sierksdorf, await discovery near the German Baltic Sea coast.

### Predicting blue carbon stocks across Germany (objective iv)

A fourth objective of the study was to use the identified relationships between seagrass meadow attributes and environmental parameters on C_org_ to extrapolate stocks for the entire German Baltic Sea region. This regional variation in C_org_ was best explained by seawater depth, seagrass complexity, and the fraction of fine particles in the sediment, not average MOV or interactions between parameters. The latter two had a positive effect on the C_org_ content in the sediment, whereas stocks decreased with seawater depth, suggesting that the highest stocks were found in shallow locations with high seagrass complexity and the ability to accumulate fine-grained particles, regardless of MOV at the seafloor. However, these parameters do not demonstrate a clear ability to predict the regional distribution of C_org_ (Fig. 5). A combination of these three parameters, along with water motion, is consistently found to be the main biotic and abiotic driver of C_org_ in the literature, but its influence varies widely across regions and seems principally coupled to local hydrodynamic regimes. Because sediment erosion and detritus export rates are typically heavily influenced by water motion (e.g. lower wave height and exposure, fetch, and currents), sediment C_org_ content is typically lower in dynamic systems compared to more static ones (e.g. Samper-Villarreal et al. 2016, Mazarrasa et al. 2017a, Prentice et al. 2019, Novak et al. 2020, Mazarrasa et al. 2021). But in the absence of strong water movement, like in the German Baltic Sea where tides are negligible, water currents are weak, and wave heights are low, other aspects of the systems can shape C_org_ stocks. The lack of a trend observed in our data in spite of sampling extremes in hydrodynamics of our system, is a testament to this hypothesis, whereby the two sites that experienced the strongest (Teichhof, Wackerballig) and the weakest (Orth, Sierksdorf) maximum current speeds at the seafloor both had the highest C_org_ pools of all sites examined in this study. Reduced hydrodynamics may also mean lower seagrass detritus export from the system and may help explain the mainly autochthonous provenance of the C_org_ and overall high stocks in our study.

**Figure 5.**
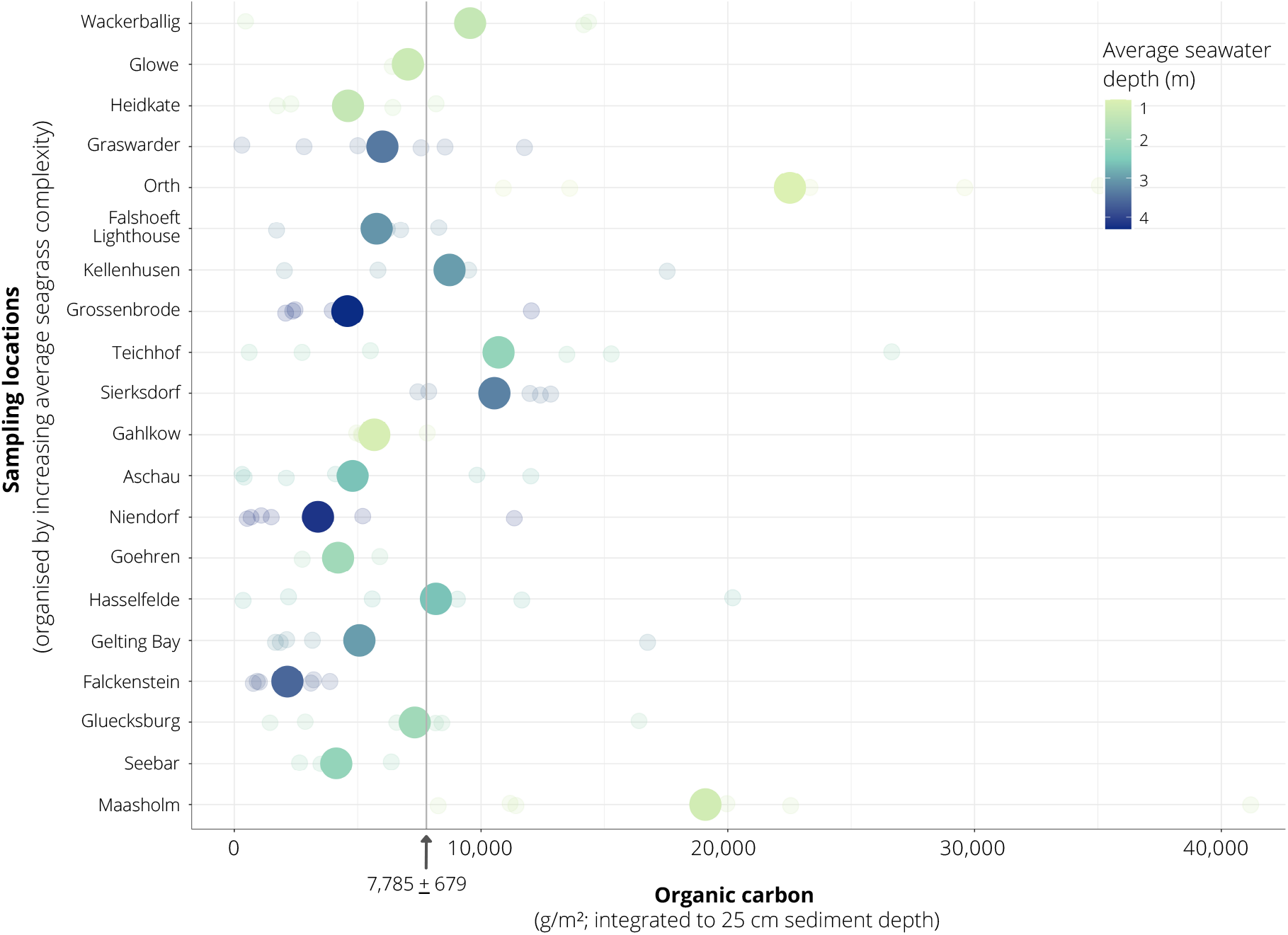
Interplay between seagrass complexity and seawater depth predictors of organic carbon (C_org_) content in seagrass-vegetated sediments along the Baltic Sea coast of Germany.

The positive relationship observed between the amount of fine particles in the sediment and C_org_ in seagrass-vegetated sediments is not surprising as it is well known that more C_org_ is associated with finer mineral particles in soils and sediments (Calvert et al., 1995; Lin et al., 2002). These fine particles nurture anoxic conditions in the uppermost layer of the soil that protect organic particles from remineralization (Duarte et al. 2013). Our findings are consistent with previous findings for *Z. marina* meadows in other than the German parts of the Baltic Sea (e.g. Sweden, Dahl et al. 2016; Denmark and Finland, Röhr et al. 2016) and elsewhere (*Z. marina* outside Baltic Sea: Miyajima et al 2015, 2017, Prentice et al. 2019, Krause et al. 2022; multiple species, including *Z. marina*: Kennedy et al. 2010; *P. oceanica*: Gacia et al. 2002, Hendriks et al 2008). For example, in Denmark and Finland, more than 40% of the variation in C_org_ between sites could be explained by sediment characteristics, including the fine sediment content.

The thick canopy of seagrass leaves is known to effectively intercept particles in the water column, as well as decrease sediment erosion and seagrass detritus export (Duarte et al. 2013). While seagrass complexity is a strong positive predictor of the regional differences in C_org_ in our study, and that of others (Jankowska et al 2016, Samper-Villarreal et al 2016, Serrano et al 2016), this was not the case for *Z. marina* meadows off the coast of British Columbia, in Canada, where no clear relationship (negative or positive) was observed between the two parameters (Prentice et al. 2019). Here, water motion was the strongest predictor of C_org_.

The dampening effect of seawater depth on surface water motion is correlated to the trend of increasing C_org_ content with deeper depths (Lavery et al 2013, Mazarrasa et al. 2017a). While we also observed a dampened effect of water currents (but at the seafloor, rather than surface waters) by increasing seawater depth (*ρ* = −0.43), an opposite relationship was seen between C_org_ content and depth. A fourfold decrease in C_org_ was also observed with increasing seawater depth (from 2-4 m to 6-8 m) in *P. oceanica* seagrass, in the Mediterranean Sea (Serrano et al. 2014). Regarding the Baltic, the realization of the Helsinki Commission’s Baltic Sea Action Plan (BSAP) goals, which implicate a considerable nutrient abatement, would result in seagrass expansion into deeper waters, as was shown by Bobsien et al. 2021 for the German part of the Baltic Sea. This expansion would lead to an enhancement of the CO_2_ storage potential by these habitats, but it is important to consider that this potential is diminished at deeper depths (relative shallow depths) - an important consideration for carbon accounting when including seagrass blue carbon contributions to the German national CO_2_ budget.

### Scaling up for CO_2_ accounting (objective v) and further considerations

Our measurements confirm that seagrass meadows in the Baltic Sea coast of Germany store a large C_org_ pool. The high spatial heterogeneity seen across the region warrant site-specific investigations to obtain accurate estimates of blue carbon. However, localities with high seagrass complexity, high fine sediment fraction, and low seawater depth could help select localities with more favorable C_org_ accumulation potential. An unexpected and significant relic terrestrial C_org_ pool was found beneath the seagrass meadow in Sierksdorf (Luebeck Bay). It is likely that many more submarine peatlands await discovery along the southwestern Baltic Sea region, and that they hold a millennial timescale C_org_ deposit similar to those found in Germany.

Based on a conservative scaling up of measurements (integrated to 25 cm sediment depth), collectively (total of approx. 285 km^2^, Schubert et al. 2015) seagrass meadows in the German Baltic Sea are preventing 8.14 Mt of future CO_2_ emissions from being released into the atmosphere. However, it must be noted that in some locations (Gelting Bay, Teichhof) the sediment has been eroded to the marl layer, which seagrass roots cannot penetrate, while in other locations the sediment is known to extend to 7 m sediment thickness below the seafloor, e.g. within the inner areas of Mecklenburg Bay (Lemke 1998), such as Sierksdorf, Niendorf, Kellenhusen, Grossenbrode in the present study.

Because C_org_ originated from within the meadow (not imported from outside its boundaries), accumulation of blue carbon in Germany may be contingent on healthy seagrass habitats. Furthermore, loss of these habitats will have negative consequences for the German remaining CO_2_ budget because the C_org_ stored beneath meadows may be rereleased into the water column and later to the atmosphere. Their loss would also impact their many co-benefits (see Heckwolf et al. 2021). Given the pressing need to offset and prevent future CO_2_ emissions, more stringent and concerted efforts are urgently needed to enhance the C_org_ storage potential (via habitat restoration and improving growing conditions) and prevent further degradation (via conservation) of seagrass habitats along the Baltic Sea coast of Germany. Our study provides urgently needed knowledge and constitutes a further incentive to enhance efforts in protecting existing seagrass meadows in Germany, and restore them where natural recolonization is likely slow, such as in enclosed embayments or areas that have seen a loss in seagrass habitat, especially where the distance to the next vegetated site is high.

## Author contributions

Conceptualization: AS, PRS, TBHR; Methodology: AS, TCÓC, PRS, WH, TBHR; Data acquisition and analyses: AS, TCÓC, PRS, WH; Writing (original draft): AS; Supervision: TBHR; Funding acquisition: PRS, TBHR. All authors contributed intellectual input, edited, and approved this manuscript.

## Acknowledgments

Many thanks are due to Ainara Zander, Christian Howe, Dr. Florian Huber, Marlene Beer, Nasif Bin Said, Philipp Suessle, Roxanna Timm for their help in the field and/or lab. Dr. Christian Hamann, Dr. Jan Dierking, and Dr. Tomas Hansen for their insights on the stable isotope analyses.

